# *StateHub-StatePaintR*: rapid and reproducible chromatin state evaluation for custom genome annotation

**DOI:** 10.1101/127720

**Authors:** Simon G. Coetzee, Zachary Ramjan, Huy Q. Dinh, Benjamin P. Berman, Dennis J. Hazelett

## Abstract

Genome annotation is critical to understand the function of disease variants, especially for clinical applications. To meet this need there are segmentations available from public consortia reflecting varying unsupervised approaches to functional annotation based on epigenetics data, but there remains a need for transparent, reproducible, and easily interpreted genomic maps of the functional biology of chromatin. We introduce a new methodological framework for defining a combinatorial epigenomic model of chromatin state on a web database, *StateHub*. In addition, we created an annotation tool for bioconductor, *StatePaintR*, which accesses these models and uses them to rapidly (on the order of seconds) produce chromatin state segmentations in standard genome browser formats. Annotations are fully documented with change history and versioning, authorship information, and original source files. *StatePaintR* calculates ranks for each state from next-gen sequencing peak statistics, facilitating variant prioritization, enrichment testing, and other types of quantitative analysis. *StateHub* hosts annotation tracks for major public consortia as a resource, and allows users to submit their own alternative models.

## Introduction

Chromatin segmentations are increasingly important for a broad area of research that includes regulatory genomics, genetic epidemiology, precision health, and molecular genetics. There is a need for consistent, unbiased resolution of chromatin states to interpret the epigenome and predict function across different tissues and cell types.

Complex, overlapping patterns of post-translational modifications (PTM) to histone subunits (Rando, 2013; Gardner et al., 2011), signify differing states of chromatin activity. These modifications consist of mono-, di-, or tri-methylation and acetylation of histone 3 lysines 4, 9, 27, and 36 (Rothbart and Strahl, 2014). Direct assays for histone PTMs with next-generation sequencing (NGS) using chromatin immunoprecipitation (ChIP-seq) result in a set of genomic intervals with evidence for enrichment over background (input chromatin), using signal intensity.

In addition to ChIP-seq of histone PTMs, there are also NGS methods for histone displacement, including DNase I hypersensitivity (Boyle et al., 2008) (DNase-seq or DHS), Formaldehyde Assisted Isolation of Regulatory Elements (Simon et al., 2012) (FAIRE-seq), Assay for Transposase Accessible Chromatin (Buenrostro et al., 2013) (ATAC-seq) and Nucleosome Occupancy and Methylome sequencing (NOMe-seq) (Gal-Yam et al., 2006). Histone displacement, nucleosome positioning and DNA methylation are also detected in genomewide assays (*e.g*. whole genome bisulfite sequencing (Cokus et al., 2008)). Histone displacement is associated with transcription factor binding and transcriptional activity (Thurman et al., 2012). In addition, direct binding of transcription factors is measured in ChIP-seq experiments with an antibody directed against a transcription factor or an epitope-tagged version.

All these data are compatible with data represented as genomic intervals (in bed format), including CpG islands, annotated transcription start sites, repeat elements, 3′ UTRs. The input and final (output) processed data format are both as browser extensible data (.bed), a flexible standard for different peak calling methods (*e.g*. “narrowPeak” and “broadPeak” are types of ․bed files).

Several machine-learning approaches integrate NGS experiments into annotation tracks (Li et al., 2015). The goal is to discover epigenomic states and aid in understanding “non-coding” genomic elements in an unbiased and biologically meaningful way. Newly discovered states are likely an amalgam of true functional categories reflected in chromatin biology. The most popular and widely used of these machine learning methods is ChromHMM (Ernst and Kellis, 2012). Other machine-learning approaches include spectral-based learning (Song and Chen, 2015), inference based on read counts (Mammana and Chung, 2015), dynamic bayesian networks (Hoffman et al., 2012), probabilistic approaches (Hon et al., 2008), supervised enhancer detection (Santoni, 2012), and other hidden markov methods (Zacher et al., 2014; Sohn et al., 2015; Biesinger et al., 2013).

The interpretability and general usefulness of the state predictions produced by these algorithms varies. A multitude of states often must be consolidated into simpler, biologically meaningful categories. Hoffman *et al.*, recognized this problem when they proposed a combined meta-analysis of ChromHMM and Segway annotations(Hoffman et al., 2013). However, a software framework for expert or rule-based segmentations is still lacking. Such methodology is needed for integrating different experimental data (including non-NGS data) in a reproducible way, reflecting both the novel insights gained from the machine learning methods and our current understanding of genome biology.

Here we introduce *StateHub* and *StatePaintR* for generating and documenting chromatin state and other genome segmentation models in a transparent and reproducible fashion. *StateHub* is a public resource for storing annotation models, state definitions and associated data in a shareable, referenceable form. The *StatePaintR* package implements these models and state definitions to produce annotation tracks. We show that *StatePaintR* can be used to rapidly annotate large collections of public data for summarizing epigenomics data or annotation of variants. We show how annotations gracefully degrade, in that cell types or tissues with missing data types are annotated appropriately based upon available information. We show some use cases, and describe how *StatePaintR* uses ChIP-seq data peak statistics to rank the state prediction for each segment.

## MATERIALS AND METHODS

### Implementation and availability

*StatePaintR* is implemented as a software package in the R language freely available from the Bioconductor repository: www.bioconductor.org/packages/release/bioc/html/-StatePaintR.html. The package contains functions for generating annotation tracks according to the rules specified in a model matrix. The matrix cell values can take any of 4 different values (Table 1) representing the relationship between the row labels (proposed state) and column labels (functional categories). The current package only accepts these 4 values, since each value produces very specific behavior from the software. *StatePaintR* includes a peak score for each state drawn from all experiment categories (columns) that have a matrix value of 3, *i.e*. because they are required for and consistent with that state. The peak scores are rank normalized on a scale of 1 to 1,000, with 1 being the minimum peak size and 1000 being the maximum. If multiple categories are required, *StatePaintR*selects the median peak score for the annotation. This behavior can be overridden (see documentation for details). The package includes an R-markdown vignette. The current release version of this vignette is always available from the Bioconductor website.

**Table 1.** *StatePaintR* matrix values. *StatePaintR* assigns annotations according to custom rules specified in a matrix. The rules are represented as an integer code that takes any of 4 values [0-3]. The meaning of each value is summarized in the table below.

| required for state? | consistent with state? | binary value | decimal value |
| --- | --- | --- | --- |
| No | No | 00<sub>2</sub> | 0 |
| No | Yes | 01<sub>2</sub> | 1 |
| Yes | No | 10<sub>2</sub> | 2 |
| Yes | Yes | 11<sub>2</sub> | 3 |

*StateHub* is implemented as an interactive website (www.statehub.org). *StateHub* contains a database implemented in [MongoDB](https://www.mongodb.com/) and a search engine written with Google Web Toolkit (GWT), which updates dynamically with user input. This database includes all models, model metadata and pre-computed *StatePaintR* browser tracks. Models are composite [JSON](https://www.json.org) objects that include an unique identifier, name, revision number, a searchable text description, and a model matrix (as defined in Table 1). The website also includes links to this manuscript, R-markdown containing code for figures, the latest version of the vignette, links to twitter feed and additional instructional materials.

### *StateHub* models

The main text makes reference to two models in *StateHub* (statehub.org). The unique identifiers of these models are as follows: “Default” (model ID: 581ff9f246e0fb06b4b6b178) and “Focused Poised promoter” (model ID: 5813b67f46e0fb06b493ceb0). In each of the two models presented and discussed in this paper we chose a naming convention for our states reflecting biological function.

### Annotation of public datasets

Preprocessed peak calls were obtained from the IHEC and ENCODE websites (see Table 2) for hg19, and where possible hg38. Where possible we used IDR (Irreproducible discovery rate) processed narrowPeak calls for DHS and broadPeaks for broad marks (H3K27Ac, H3K4me1, H3K27me3, H3K36me3) unless otherwise specified in the model. A complete manifest with filenames, plus all annotation tracks are available on the *StateHub* website.

### Enrichment calculations

*Parkinson’s GWAS variants*. To illustrate the use of StatePaintR chromatin state segmentations in GWAS functional annotations, we revisited an earlier study of Parkinson’s disease in which tested for tissue-specific enrichment of genetic associations. Parkinson’s GWAS variants were obtained from a previously published large scale meta-analysis (Nalls et al., 2014). We used a beta-binomial conjugate distribution to estimate the credible range of differences in overlaps between observed (GWAS hits) vs. random variants. To calculate enrichment we selected all variants within 1 MB of the index SNP in each region with a minor allele frequency (MAF) > 0.01, defining foreground as SNPs in linkage disequilibrium with the index SNP at a cutoff of *r*^2^ > 0.8 and background as all SNPs inclusive (MAF > 0.01). *Enrichment in genomic annotations*. Analyses and graphics were produced using the SegTools package (Buske et al., 2011).

### Analysis of methylation data

To select methylation variants, we analyzed the Infinium HM450 data of 114 ovarian tumor samples (Patch et al., 2015) and 216 control normal Fallopian tube samples (Teschendorff et al., 2016). We define differentially methylated regions as those having a difference in beta values of 0.3 (cancer vs. normal) and significance in Mann-Whitney U-test (FDR-corrected p-value < 0.01). We then performed enrichment calculations using overlaps between probes that were hyper-methylated in cancer *vs*. normal and the state calls from two models described above and in the text. The enrichment calculations were as described in the previous section, treating methylation “variants” analogously to SNPs. We used the complete HM450 probeset as background.

### Code for figures and tables

All code used to generate figures, tables, and this manuscript is included as an R-markdown document (additional file 1). A copy of this document may also be obtained from the *StateHub* website.

## RESULTS

### A framework for rules-based annotation

In order to assign chromatin states, it is necessary to account for the complex interplay of input from genomic annotations and cell-type-specific experimental data sources that define and demarcate functional regions of the genome (Rando, 2013). Computationally they have to be put in the right order to avoid erroneous overwriting of information-rich categories with information-poor ones.

We initially wrote a model as a decision tree, encompassing a set of basic rules for annotation, but quickly discovered that any small change to the model necessitated a near complete re-write of our software. Secondary to this, we wanted a solution that would enable us to specify any change in the model and have it produced the same way as all previous models while minimizing software updates. And thirdly, we felt that any such model should be reproducible, documented, cite-able and extensible to any combination of experiments. Moreover from a bioinformatics perspective, we felt that any two colleagues working separately should be able to produce precisely the same annotations from the same datasets and models. To satisfy these different requirements we separated the model specification from the annotation tool. We implemented model-specification as a decision matrix, which has the advantage of separating model specification from software, enabling complete explicit control of the annotation software without computer programming expertise.

We created a searchable website, *StateHub*, to host a permanent repository of models, document model objects and make them available as a resource to the community. The *StatePaintR* package retrieves models from *StateHub* and performs annotations on local data. Thus, *StateHub-StatePaintR* is a framework to document models and apply them to annotate genomic data. The models in *StateHub* consist of an abstraction layer, defining the relationships between data sources and functional categories. These categories are integrated to produce annotations (left hand column, “Chromatin States”) via a decision matrix (figure 1). Within the model each state has associated descriptions of arbitrary length which may contain key words or other relevant details (bottom right).

**Figure 1.**
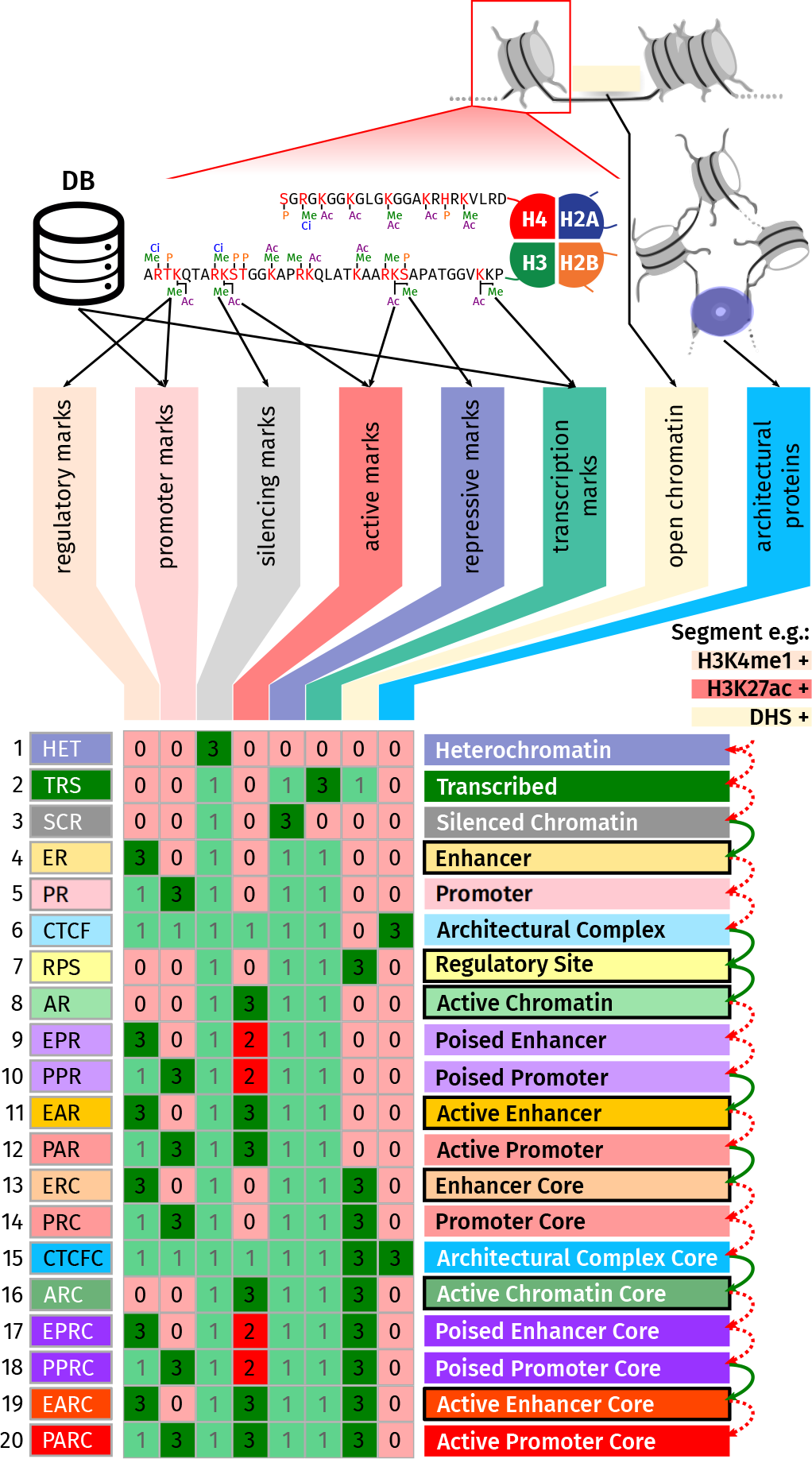
Mapping datasets to functional significance annotations. Experimental data and external database annotations are combined into abstraction layers (columns), integrated to produce chromatin states (rows) from the decision matrix. *StatePaintR* produces state assignments by iteratively comparing the marks that are present in each segment with each row of data in the table. The values of color-coded squares signify relationship between data and state: 0 (light red) the feature/data type negates the state but is not required to be present, 1 (light green) feature is consistent with the state but not required, 2 (red) if the feature is required to be available and negates the state, and 3 (green) it is both required and consistent with the state. Information content (sum of row values) of states increases from top to bottom. For the example, red dotted arrows indicate non-matching rows, and green arrows indicate matching rows. The state call corresponds to the last matched row.

Each cell of the decision matrix relates functional category to chromatin state in a 3 bit code representing the answers to two TRUE/FALSE questions (see Table 1). Is the functional category required in order to call the state? And, is overlap consistent with the state? For example, in our focused poised promoter model, the cell of the decision matrix defining the relationship between the state “PPR” and the functional category “PolycombNarrow” is 3, representing the binary value 11_2_. In order to call the PPR state, PolycombNarrow data is required be present, and second, it must also overlap with a peak. A score of 2 representing the binary value 10_2_, as in the category “Active” and state PPR, indicates that data relating to the active mark must be present to call PPR, but must not overlap. A score of 0 representing the binary value 00_2_, as in the cell for category “Core” and PPR, indicates that it is not necessary for data consistent with Core marks to be present. If the data is present and overlapping a segment, the state is excluded. The category “Translation marks” does not affect PPR in this model, even if it overlaps. Marks that are essentially irrelevant to PPR such as this one are assigned 1 representing binary 01_2_.

Thus, each row (as “state”) has a unique combination of matrix values, and the rows are organized by the software in order of increasing information content (as row sums). *StatePaintR* first generates a GRanges list (an R object containing a list of chromosomes and interval coordinates with arbitrary metadata columns attached) of all uniquely mapping segment boundaries from the start and end coordinates of every peak in all files. *StatePaintR* then evaluates the presence or absence of each mark and eliminates erroneous states. Next the program assesses overlaps of each segment to determine whether the conditions specified in each cell of the decision matrix are compatible with that segment, producing a boolean value. Rows with perfect matches in all cells are candidate state calls. Since *StatePaintR* evaluates in order of increasing information content, lower information states can be overwritten if higher information states match. This is very useful for building degeneracy in a model. An example of this in Figure 1 is illustrated by the states ER and EAR. If active marks (*e.g*. H3K27Ac) are not available for a given cell type, StatePaintR will annotate H3K4me1 marks as ER under our default model. In a different cell type for which H3K27Ac data are available, *StatePaintR* will know to distinguish between H3K4me1 enriched regions as either active or poised based on overlap of this second mark. Thus, a model can specify different state calls as appropriate based on the availability of data for each cell type.

### Segmentation of public datasets

We generated annotations of 119 ENCODE cell lines (Dunham et al., 2012), 128 Roadmap tissues (Roadmap Epigenomics Consortium et al., 2015), 26 cell lines and tissues from CEEHRC (peak calls obtained from the IHEC website), and 23 blood cell types from Blueprint (download at statehub.org). On a desktop PC it takes approximately 12-15 seconds to produce an annotation from a typical cell line, depending on the number of datasets and intervals (see S1 Figure). *StatePaintR* produces genome-browser compatible BED files with color-coded state annotations (specified in *StateHub* model). Figure 2 shows a representative region around the *POLR2A* gene from a subset of 77 high-quality(at least 15m reads) tissue samples and cell lines with H3K27Ac data from Roadmap. A complete manifest for processing these data is included in additional files 1.

**Figure 2.**
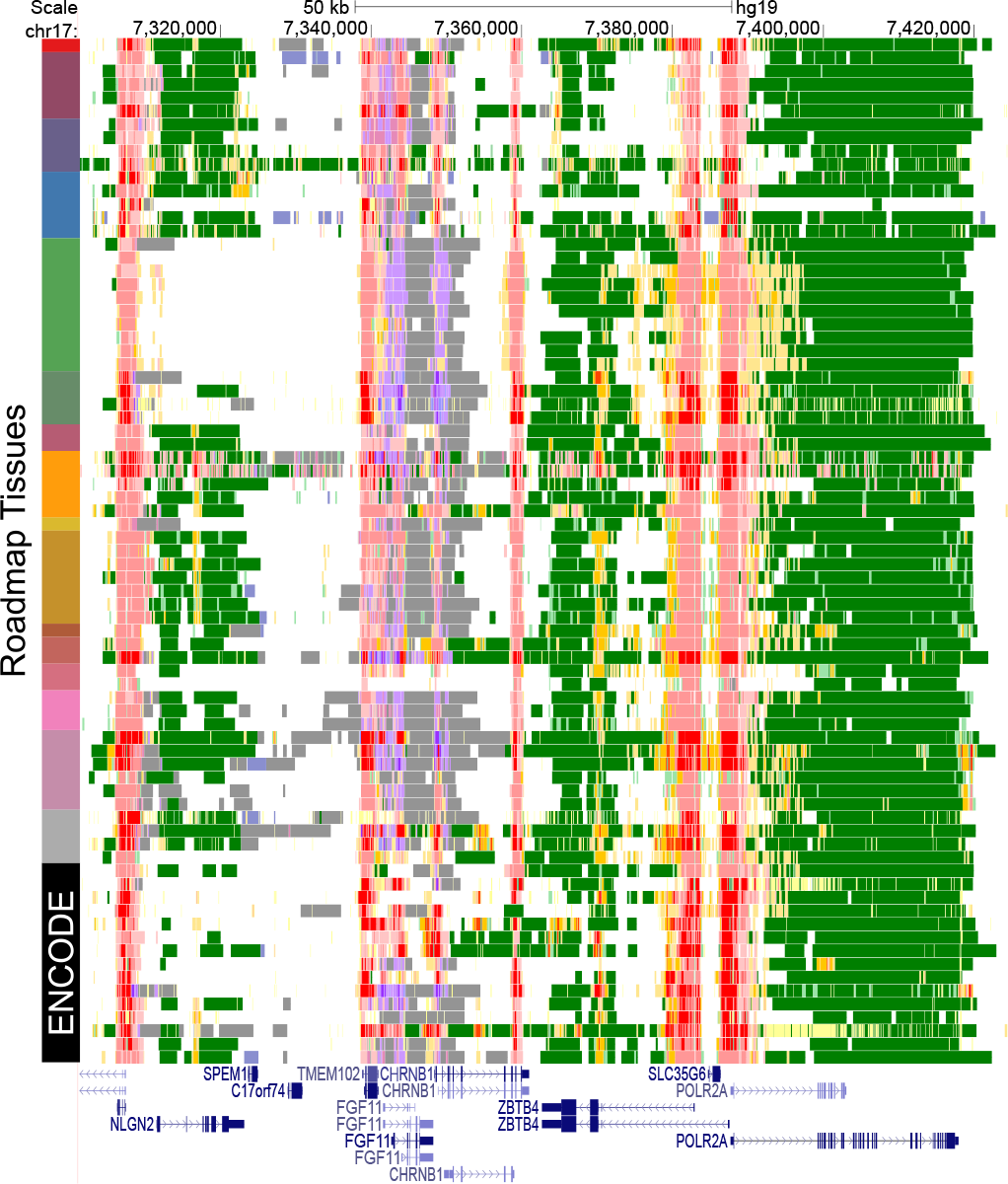
Annotation of public epigenomics data sets. Annotations of 77 cell types from the Roadmap Epigenomics consortium, including some Roadmap-processed ENCODE data, selected for their high quality with default model. Roadmap tissues are clustered and color coded at left according to the same color scheme used in Roadmap publications (Roadmap Epigenomics Consortium et al., 2015).

**Table 2.**
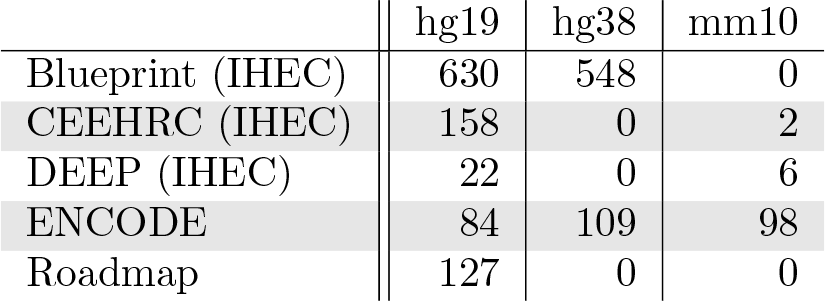
Annotation of public data sets. Data from the indicated public consortia were downloaded and processed in *StatePaintR*. The resulting annotation files and browser sessions are available from the StateHub web page under each model page.

### Annotation of genomewide association studies

A common use of genome annotation is to assign putative function to genetic loci identified by genome-wide association studies (GWAS), particularly for non-coding regions. We previously used a custom annotation of Roadmap tissues based on the approach described in this manuscript to identify locus-specific tissue enrichment in variants associated with Parkinson’s disease (Coetzee et al., 2016). In that study we displayed locus-by-tissue enrichment as a heat-map. Here we present a similar analysis using our new *StateHub* model as the basis for an alternative visualization. Since we showed that Parkinson’s disease variants are primarily associated with enhancers and promoters (Coetzee et al., 2016), we plotted the 95% range of credible values for enrichment in enhancers and promoters vs background SNPs (matched for GC content & minor allele frequency). Each locus (row) is plotted against a selection of tissues in Roadmap (figure 3).

**Figure 3.**
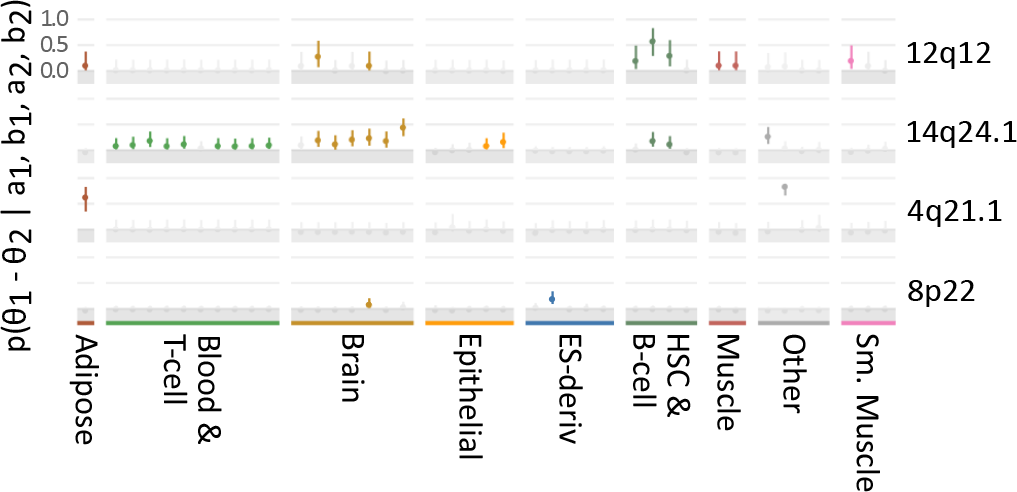
Locus- and tissue-specific enrichment of Parkinson’s GWAS variants. GWAS data and LD proxies ≥ 0.8. Bars: 95% credible range for enrichment in active states vs SNPs in the region with similar minor allele frequency and LD < 0.8, for each of 4 independent genetic loci. *θ*_1_, *θ*_2_ relative enrichment in foreground and background sets, respectively. *a*_1_, *b*_1_ number of foreground SNPs overlapping biofeatures or not-overlapping, respectively. *a*_2_, *b*_2_ number of background SNPs overlapping biofeatures or not-overlapping, respectively. a and b are shape parameters of a beta distributed prior. Significant enrichment profiles for roadmap tissues are displayed in color (REMC lineage-specific colors); non-significant are gray.

### Evaluation of two models with respect to cancer methylation

Our “default” model proposes a class of enhancers and promoters in a poised state (EPR and PPR). These features have H3K4me1 or H3K4me3 and lack H3K27Ac. This model also classifies H3K27me3 as silenced/polycomb repressed (SCR). To investigate functional enrichment of methylation variants, we looked at how differentially methylated regions (DMR) in ovarian cancer tumors partition between chromatin states as defined in this model (figure 1).

From previous work CpG islands containing temporarily silenced (poised) genes by polycomb repressive complex in normal tissues may acquire DNA methylation during cancer formation resulting in permanent silencing (Gal-Yam et al., 2008; Teschendorff et al., 2016). While the segments called EPR and PPR were associated with hypermethylated probes in ovarian cancer across tissues, the magnitude of enrichment was not great (see figure 4, “Model 1”), and it remained possible that our state definitions were too broad.

One hypothesis is that poised promoters are distinguishable by the presence or absence of focused H3K27me3, in particular the narrowPeak calls (as opposed to broad, low-level enrichment from broadPeak files used in model 1). To address this hypothesis, we repeated the analysis in figure 4 for an alternative model (model 2; “focused poised promoter”) in which H3K27me3 is called as both broadPeak and narrowPeaks. We use the H3K27me3 broadPeak file as in the previous model to identify repressed regions, and H3K27me3 narrowPeaks to identify poised states (EPR and PPR). Enhancers lacking H3K27Ac and H3K27me3 were classified as weak enhancers and promoters (EWR and PWR, not shown in figure 4). Regulatory elements with these properties have been also been called “primed” (Calo and Wysocka, 2013).

We found greater enrichment when we defined poised states in this way (compare model 2 (focused poised promoter) with model 1 (default) in figure 4). The hypermethylated ovarian cancer CpGs were more enriched in EPR, PPR, and SCR states as defined in the focused poised promoter model relative to the default model, and hypomethylated probes were enriched only in HET and SCR states (not shown). The odds ratio of enrichment for hypermethylated CpGs in EPR and PPR from the default model fell in a range between 0 and 5. However, the enrichment of the hypermethylated probes in our focused poised promoter model was > 5 in PPR and > 10 in EPR (figure 4, model 2). Thus, ovarian hypermethylated probes are enriched across roadmap tissues in H3K27me3+ enhancers and promoters, and we concluded that H3K27me3 narrowPeaks are an important distinguishing feature for this class.

**Figure 4.**
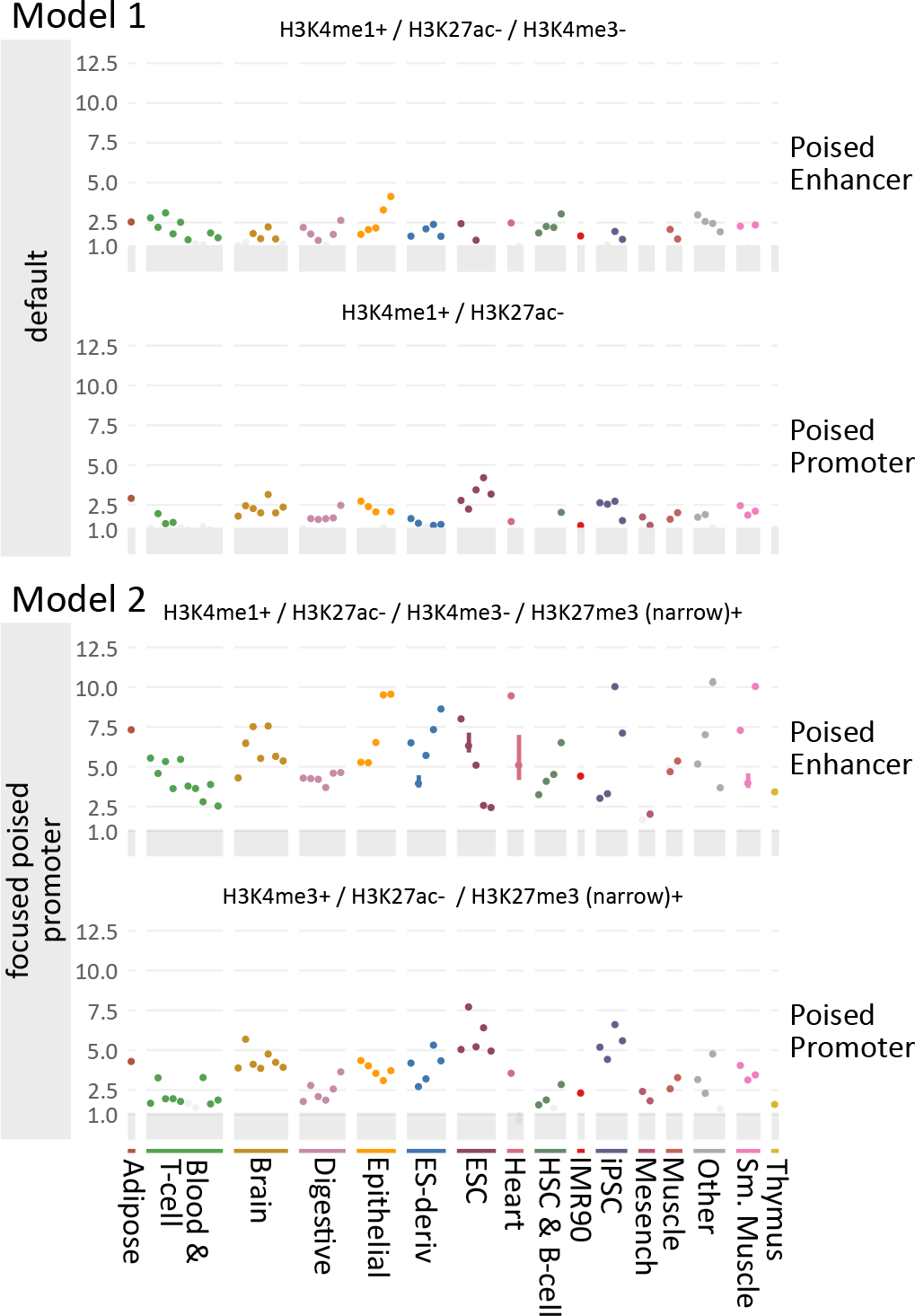
Example of model comparisons. Enrichment as in figure 3 using either of two different state models (model 1 and model 2) from StateHub, “Default” and “Focused Poised Promoter”, which differ in the treatment of poised promoters. Y-axis range is the same for both plots. Both models distinguish hypermethylated probes in the poised state but model 2 is more selective than model 1. In this model (2) enhancers with H3K4me1 and promoters with H3K4me3 overlapping narrow regions of H3K27me3 are poised (EPR and PPR), but those without H3K27me3 are called weak (EWR and PWR). Model 1, by contrast, assigns promoters lacking active marks to the poised state.

### Enrichment of functional annotation

Next, we characterized the distribution of states in our focused poised promoter model relative to Gencode v37 gene annotations and also to enhancers from Ensembl (Zerbino et al., 2015). Figure 5 shows the relative enrichment of Human mammary epithelial cell (HMEC) chromatin states in each of these features. We found enrichment in Ensembl enhancers for three states: Active enhancer (EAR), Active regions (AR) and Weak enhancer (EWR). The definition of “active enhancer” in the Ensembl build is cumulative across cell types (Zerbino et al., 2015) and therefore includes many cell-type specific enhancers that would be predicted to be weak (having exclusively H3K4me1) in a particular cell line such as HMEC. These three states were not enriched in any other category of genomic annotations. Likewise we found enrichment of the inactive enhancers in Transcribed (TRS) and Silenced/Polycomb (SCR). TRS was most enriched in gene body annotations, particularly internal exons and introns. SCR and Heterochromatin (HET) were depleted across all categories. Lastly, the 5′, first exon and first intron regions were enriched in active and weak promoters, consistent with the role of these regions in transcription initiation.

**Figure 5.**
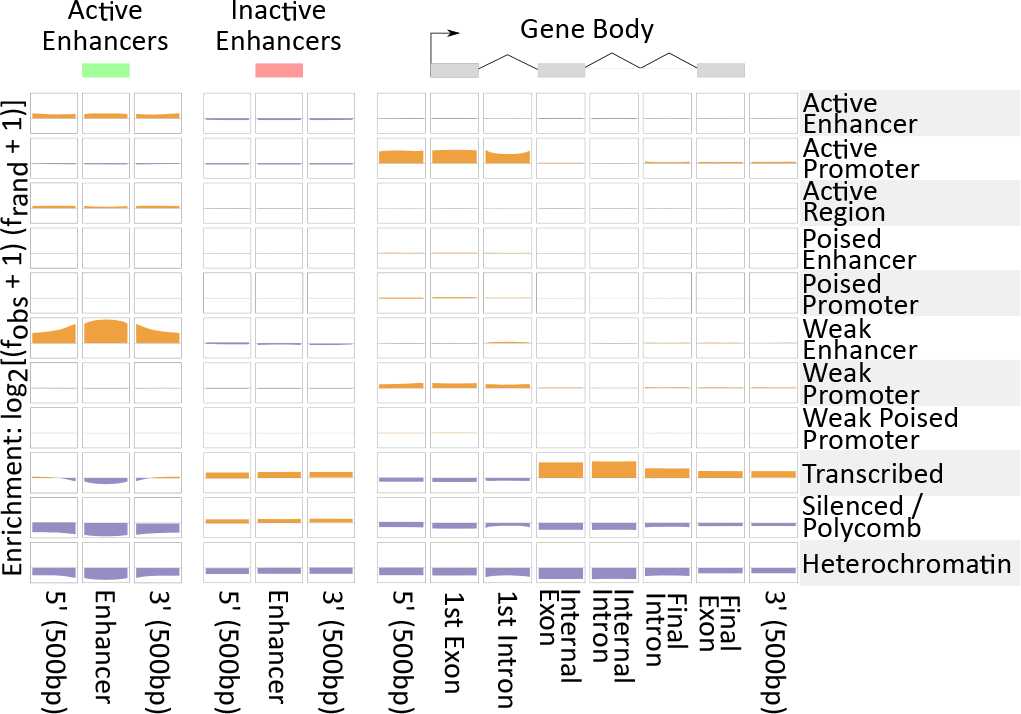
Enrichment in genomic annotations. Relative enrichment of called states genomewide from HMEC in annotations from Ensembl and Gencode. Genegraph (top) visualization of the regions indicated for each column. Enrichment is *log*_2_ observed over random. Positive enrichment is indicated with mustard color (scale from 0 to 0.66) *vs*. relative depletion in purple (scale from 0 to −0.37).

### Enhancer predictions

To use ChIP-seq data for quantitative analysis, we ranked within each state by peak score from Macs2 output (generic peak height). We programmed *StatePaintR* to rank each state by normalizing on a scale of 1-1000, 1000 being the highest rank. *StatePaintR* ranks the required dataset(s) for each state (*i.e*. assigned “3” in the decision matrix). To evaluate the ranking function, we measured area under the precision-recall-gain curve (AUPRG) using the set of experimentally validated human and mouse noncoding fragments with gene enhancer activity as assessed in transgenic mice (VISTA enhancer browser and (Visel et al., 2006)). We randomly sampled 100 enhancers from 7 VISTA tissues to evaluate different aspects of our models (training), and then used the remainder of the data to test our enhancer predictions against previously published predictions using the same data sets.

**Table 3.** Relative performance of *StatePaintR* enhancer ranking *vs*. VISTA enhancers (Visel et al., 2006). Columns 2-6 reflect the area under the precision-recall gain (auprg) curve.

| source | neural tube | mid-brain | hind-brain | limb | heart | average auprg | average rank |
| --- | --- | --- | --- | --- | --- | --- | --- |
| ENCODE | 0.82 | 0.82 | 0.80 | 0.85 | 0.88 | 0.83 | 3.4 |
| EnhancerFinder | NA | 0.59 | 0.63 | 0.67 | 0.82 | 0.54 | 6.8 |
| REPTILE | 0.86 | 0.87 | 0.76 | 0.89 | 0.92 | 0.86 | 2.0 |
| DELTA | 0.81 | 0.84 | 0.76 | 0.84 | 0.93 | 0.84 | 3.6 |
| RFECS | 0.79 | 0.85 | 0.78 | 0.85 | 0.92 | 0.84 | 3.0 |
| CSIANN | 0.72 | 0.68 | 0.62 | 0.69 | 0.84 | 0.71 | 6.2 |
| StatePaintR<sup>†</sup> | 0.84 | 0.84 | 0.79 | 0.85 | 0.88 | 0.84 | 3.0 |

<sup>†</sup>Annotations using “poised promoter” model as described in the text.

Some states, including the ones that are germane for enhancer prediction, reference more than one required (matrix value 3) dataset, and therefore it was necessary to optimize the best method for ranking based on > 1 ChIP-seq experiment. We computed the average, median and ceiling functions of ranks across multiple ChIP-seq tracks. The three methods were comparable, but median and average produced the best results (S2 Figure).

There are three required marks for active enhancers in our model, but if one of them is not informative for active enhancer prediction, using the ceiling “max” method would produce false positives when this mark has the highest peak rank. Therefore we interrogated which marks are informative using a leave-one-out approach. We found that leaving out H3K4me1 significantly improved our predictions, whereas leaving out the other marks did not (S3 Figure).

Next we assessed AUPRG of different state calls vs VISTA enhancers and found that predictive power descends in order AR + EAR > EAR > AR > RPS > EPRC > etc (S4 Figure). When we tried combinations of states the highest precision recall gain was observed for EAR, EARC, AR and ARC added together (S4 Figure), and this was greater than other combinations and than any of the state calls individually. H3K27Ac is the only mark common to all these states, suggesting that H3K27Ac is the most informative predictor of enhancers.

Since H3K4me1 does not improve predictions and is the only thing that distinguishes between AR and EAR (by its presence or absence), an improved model would consolidate AR and EAR into a single state and reassign “1” to H3K4me1 instead of “3”, leaving this mark exclusively to define weak (or primed) promoters.

To validate our method of enhancer prediction, we compared our predictions with ENCODE Encyclopedia, Version 3 (zlab-annotations.umassmed.edu), EnhancerFinder, RFECS, DELTA, CSIANN, and REPTILE (Erwin et al., 2014; Rajagopal et al., 2013; Lu et al., 2015; Firpi et al., 2010; He et al., 2017) for held-out data using AUPRG (S5 Figure) (Flach and Kull, 2015).

Our predictions are comparable to the Encode model which uses H3K27Ac overlapping with distal DHS, RFECS and REPTILE, which had the lowest average rank across tissues (table 3, S5 Figure). Our predictions compared favorably to EnhancerFinder and CSIANN which had an average rank > 6 across the different tissues; heart, midbrain, hindbrain, neural tube and limb. Predictions are only available for these tissues. Thus, *StatePaintR* ranking is useful for drawing quantitative comparisons between different models, making predictions, or prioritizing regions for functional evidence.

## DISCUSSION

We created a platform for hosting, browsing, and generating new genome annotation models called *StateHub*. The *StateHub* framework makes it possible to specify combinations of genomic data as they relate to regions of functional significance in epigenetically marked chromatin. In addition we created a software package, *StatePaintR*, that facilitates the use of *StateHub* models to generate browser tracks for bioinformatic analyses. We showed how *StatePaintR* can be used as part of a workflow with uniformly processed data to generate reproducible annotations from public and private data.

Our framework does not replace current machine learning methods, the aim of which is to discover states. But these methods suffer from certain drawbacks that we have addressed with a rules-based approach that provides greater transparency and reproducibility. For example, it is often the case with machine-learning methods that more states are discovered than immediately understood, and there have been different solutions proposed. During discovery, one could iteratively reduce the number of states, minimizing the number of similar or redundant combinations of histone marks. Then the number of discovered states would depend on the number of unique data types used for learning and their distribution around known features. This brittleness makes replication in different settings (with different labs or types of experiments) impractical. Our method avoids these issues, allowing users to specify a model of the epigenome in a matrix (as in figure 1) that accounts for all known possibilities. Thus, we built a comprehensive framework for a rules-based annotation, reflecting current hypotheses (or models) of the epigenome.

A significant drawback of our approach is that some unusual combinations of marks that may have biological function will be ignored. This has much to do with the fact that *StatePaintR* is not for discovering novel states, but rather for annotating the genome according to a specific, existing model. Nonetheless, the label assignment step of other chromatin state discovery tools also suffers the same limitations; states are aggregated or optimized in an iterative fashion based on prior knowledge and assumptions. ENCODE for example has published tracks for both ChromHMM and Segway that include multiple states with similar names (*e.g*. “Tss” vs. “TssF” from ChromHMM, and “EnhF1” vs. “EnhF3” from Segway (Hoffman et al., 2013)). To resolve discrepancies between the two methods, the authors of those studies proposed a combined analysis to simplify the number of state labels and summarize discovery using a rule-based metric not unlike a *StateHub* model. Thus, they classified regions into 7 types “emphasizing biologically meaningful differences” (Hoffman et al., 2013). In direct comparisons we found that our own annotations exhibited greater similarity to the combined analysis than to either of the Segway or ChromHMM tracks separately (not shown). Whatever the protocol, the basic problem persists; machine-learning is able to provide insight into what the categories are, but not how many categories there should be. Currently this remains the exclusive province of the biologist.

One of the additional challenges is compatibility between data sets. In order for two or more cell types to be annotated according to the same model, it is necessary to combine each of the cell types for the training. One solution is concatenation of genomes (Hoffman et al., 2013). Another approach is to jointly model epigenomes in parallel, as proposed in Integrative and Discriminative Epigenome Annotation System (IDEAS) (Zhang et al., 2016). This approach has the distinctive advantage of also modeling segment boundaries. Our approach does not model boundaries, but does offer some advantages. One is reproducibility: *StatePaintR* always produces the same annotation independently for each cell type from the same model. Secondly, even samples with different types of data or missing data result in compatible annotations because they come from the same model. Third, the models, composed of a 2D matrix with a range of 4 values, are relatively easy to understand and author. Every file produced in *StatePaintR* contains a record of the model ID, genome version and all the source files. Clinicians working with human genetics will value consistency and reproducibility across datasets. We produced annotations for REMC, ENCODE, IHEC and blueprint and made these available on the *StateHub* website for the two models described in this paper. The website also has links to browser sessions where they can be explored and used to create figures. A fourth advantage is speed: samples can be processed in parallel and there is no computationally expensive learning step.

A final feature that is very useful is the ranking by peak score (figure S5 Figure). Using this scheme, we investigated what states contribute most to true enhancers (S2 Figure–S4 Figure). We found that H3K27Ac defined the best predictive subset of annotations for VISTA enhancers. We also investigated different approaches for handling multiple peak calls for a state and found the median to be optimal (S2 Figure), and incorporated this method as the default behavior of *StatePaintR*. When we compared our predictions to held-out data, they were comparable to the best enhancer predictions (He et al., 2017; Rajagopal et al., 2013) and ENCODE enhancers (Dunham et al., 2012) and on the web (unpublished).

We demonstrated a workflow wherein new models generate annotations, which are used to test predictions against experimental data, and then in turn to make improvements to old models. We anticipate that this will be valuable in testing new ideas and hypotheses generated from unsupervised methods. The ability to rank features also aids in prioritizing variants for GWAS and studies of somatic mutations. Knowing which variants overlap features in the epigenomic landscape of a particular cell type is key. In the future, other methods may become available for incorporation into *StatePaintR* but the models described in *StateHub* will remain stable.

## CONCLUSION

We introduced two new computational resources, an online database of chromatin state models and processed genome segmentations called *StateHub*, and an R/Bioconductor tool called *StatePaintR* which translates epigenomics files into segmentations using these models. One may annotate incomplete datasets rapidly and sensibly according to a single model specification that gracefully degrades to lesser annotations with missing data. Annotations have header documentation with genome version, *StateHub* model, and the names of source files and their mappings. These tools document segmentations and state labels precisely as they are used in individual studies and to allow comparisons between evolving models of epigenomic states as they relate to NGS experiments. They also enable mixing of epigenomic states with other types of data, such as 3D looping assays, transcription factors, primary sequence features such as position weight matrices, or disease variants.

## ACKNOWLEDGEMENTS

The study was funded by NIH UO1CA184826 (BPB) and RO1CA190182 (DJH).

## Supporting Information

The following are additional files containing manifests to run *StatePaintR* with current releases of all public datasets listed in Table 2, links to segmentation tracks, and all code used for analysis and generation figures in this manuscript.

## Additional file 1

Complete code generated from R markdown (Rnotebooks/html format) for generating all analyses, figures and tables is available here.

## Supplementary Figures

### S1 Figure

**Figure S1.**
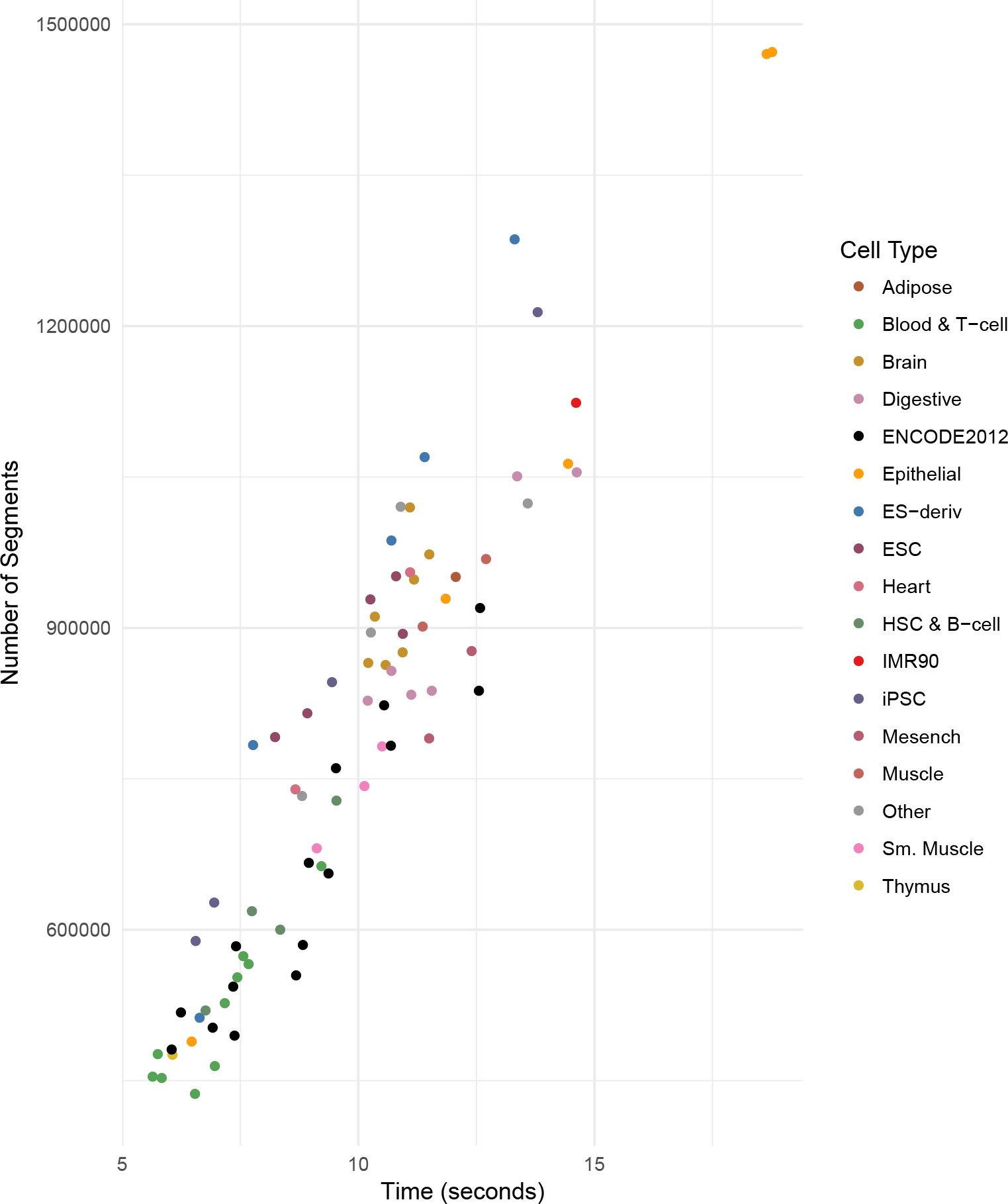
Relationship between data and runtime. *StatePaintR* takes only a few seconds to run. The exact time depends on the number number of unique segments (lines of data) created by overlapping genomic intervals of all input files, cumulative. Thus, 128 Roadmap tissues can be run in 10*s* × 128 ≈ 1, 280*s* (21*m*).

### S2 Figure

**Figure S2.**
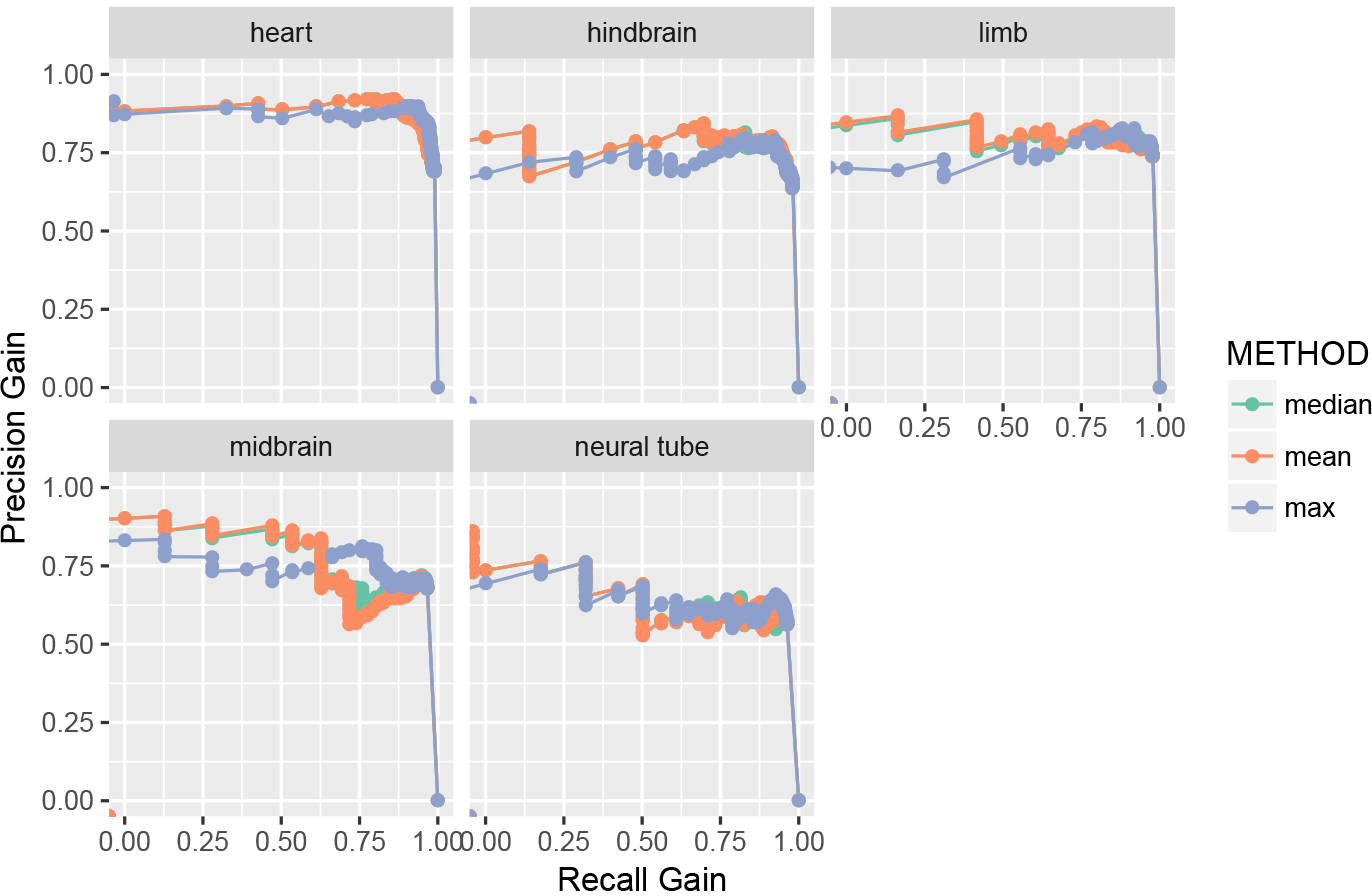
Predictions with multiple marks. Ranked ChIP-seq peak scores for multiple marks were used to rank active enhancers (H3K4me1 + H3K27Ac + DHS) by 3 methods (median, mean, ceiling) and compared to a sample (*n* = 100) of experimentally validated enhancers. The average or median of three marks was a better predictor than ceiling. The choice of function is subservient to choice of data for ranking-if one of the three is less informative, it will produce false positives when using the max method-therefore it is better to eliminate uninformative marks. See also S4 Figure.

### S3 Figure

**Figure S3.**
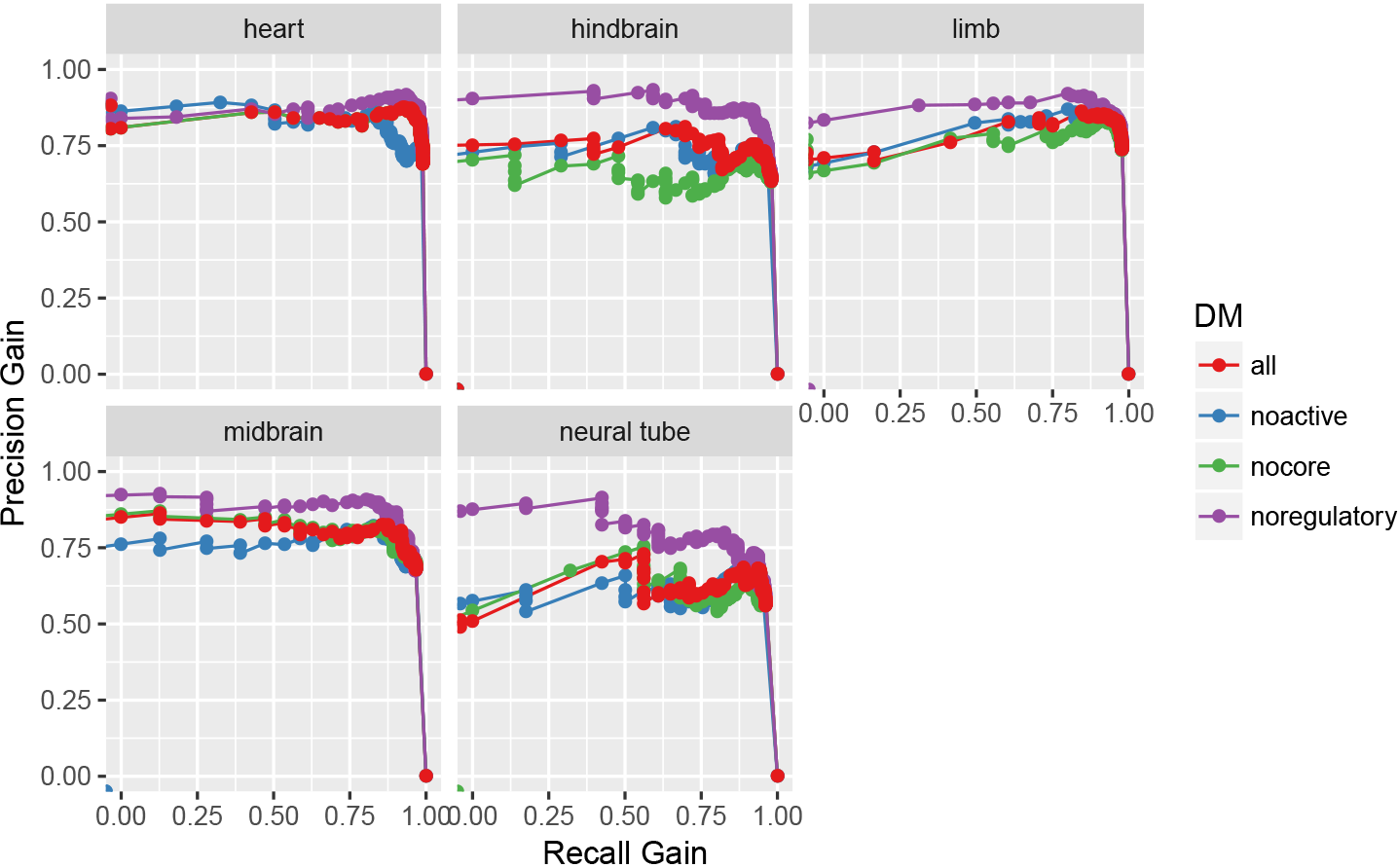
Ranking enhancers with subsets of marks. Combinations of marks were used to predict active enhancers by the max ranking method (as in S2 Figure) and compared to enhancer score. “All” includes regulatory (H3K4me1), active (H3K27Ac), and core (DHS). We also tried a leave-one-out strategy for each of these categories in succession. Leaving out H3K4me1 (“no regulatory”) produced superior predictions, suggesting that its inclusion made the predictions less specific.

### S4 Figure

**Figure S4.**
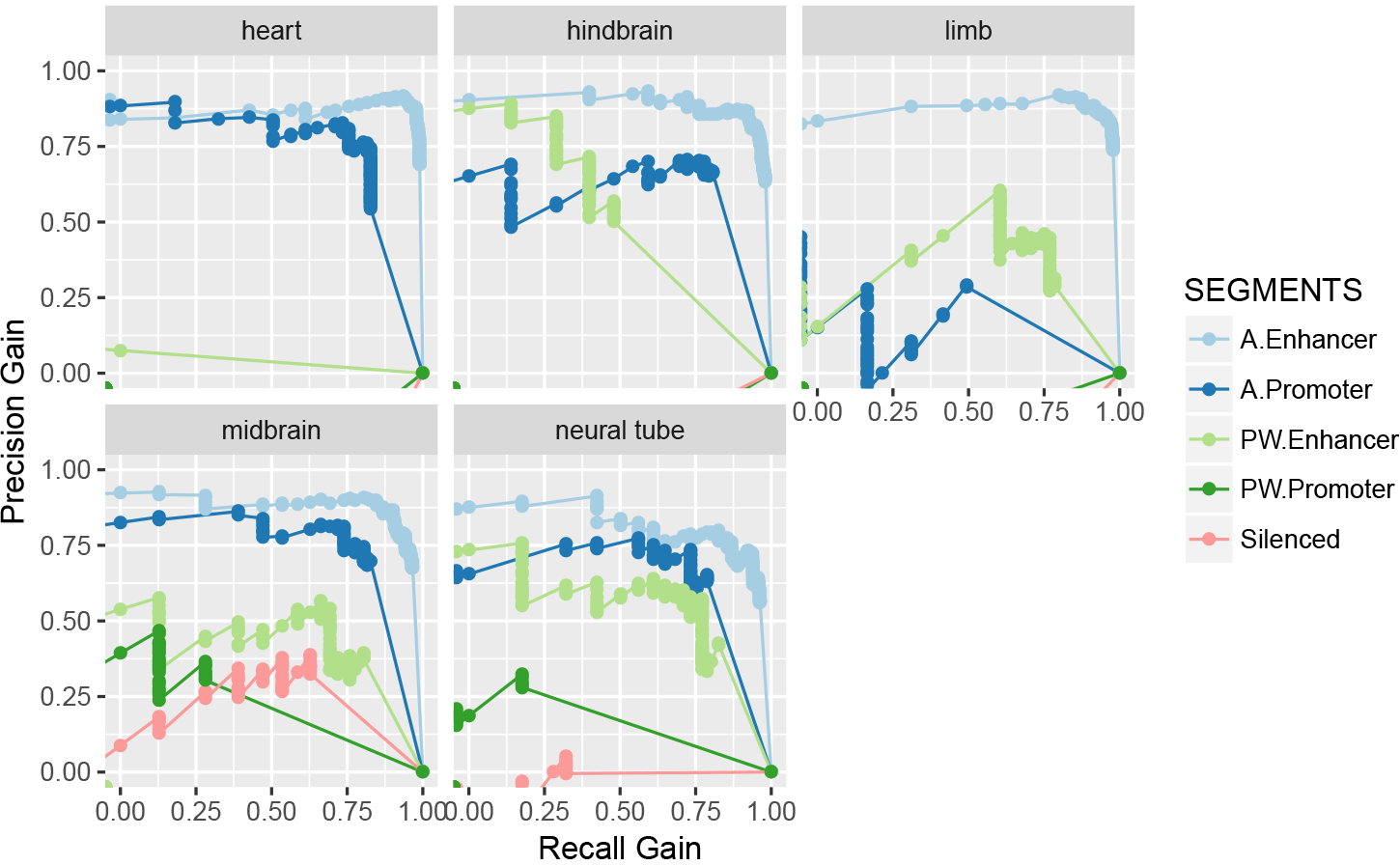
Chromatin states as predictors of true enhancers. We tested different chromatin states for their ability to predict true enhancers under the poised focused promoter model. Active enhancers exhibited the greatest predictive power under the precision recall gain curve.

### S5 Figure

**Figure S5.**
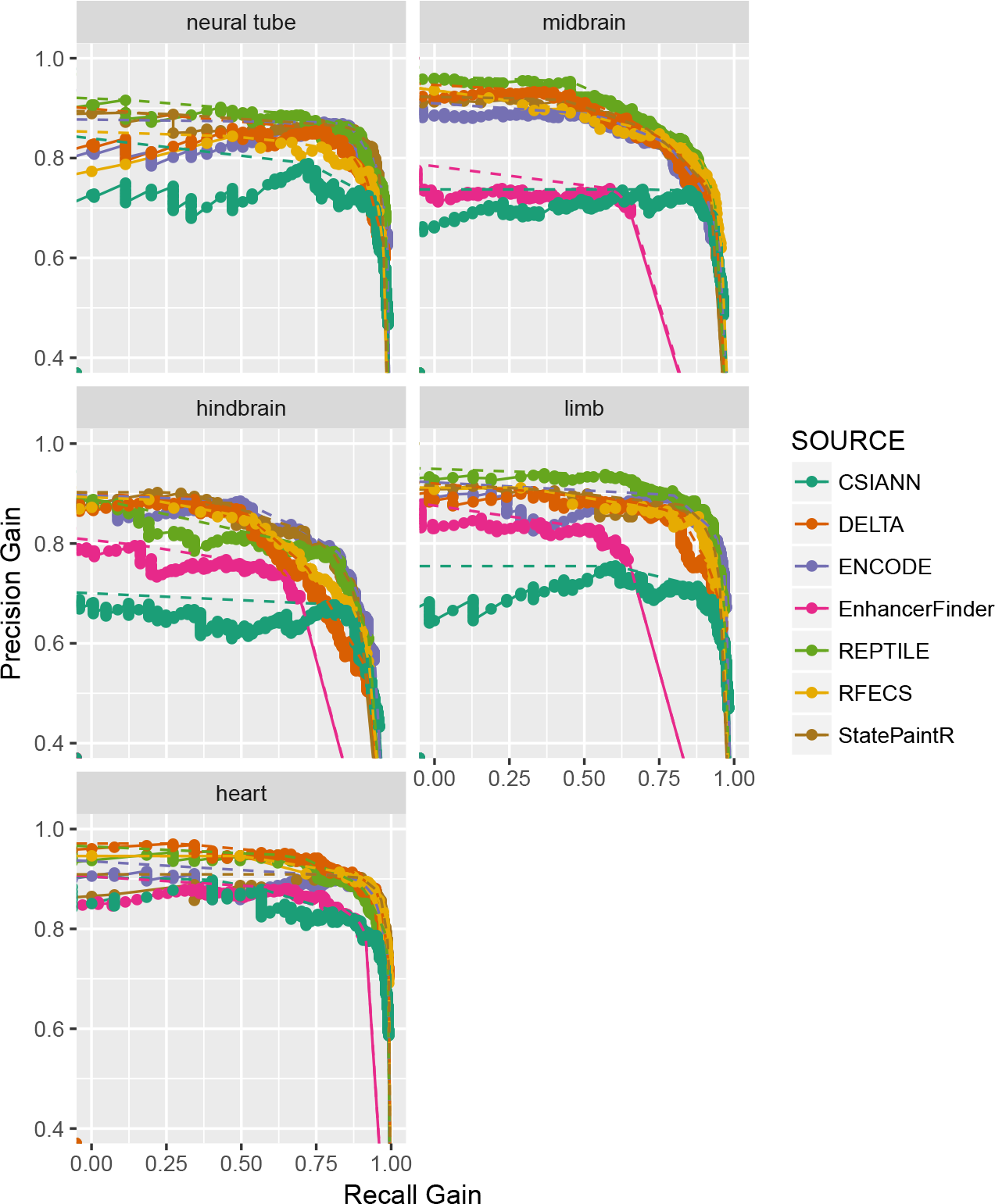
Performance of enhancer predictions. Area under precision-recall-gain curves reflect the accuracy of three models of enhancer prediction. True positive enhancers are those validated in the VISTA enhancer browser. The ENCODE method (in blue) and the *StatePaintR* method (in red) show similar accuracy in retrieving VISTA enhancers showing tissue specific enhancer activity, while EnhancerFinder (in green) is less accurate.

